# Migration of intestinal dendritic cell subsets upon intrinsic and extrinsic TLR3 stimulation

**DOI:** 10.1101/785675

**Authors:** Agnès Garcias López, Vasileios Bekiaris, Katarzyna Müller Luda, Julia Hütter, Konjit Getachew Muleta, Joy Nakawesi, Isabel Ulmert, Knut Kotarsky, Bernard Malissen, Meredith O’Keeffe, Bernhard Holzmann, William Agace, Katharina Lahl

## Abstract

Initiation of adaptive immunity to particulate antigens in lymph nodes largely depends on their presentation by migratory dendritic cells (DCs). DC subsets differ in their capacity to induce specific types of immunity, allowing subset-specific DC-targeting to influence vaccination and therapy outcomes. Faithful drug design however requires exact understanding of subset-specific versus global activation mechanisms. cDC1, the subset of DCs that excel in supporting immunity towards viruses, intracellular bacteria and tumors, express uniquely high levels of the pattern recognition receptor TLR3. Using various genetic models, we show here that both the cDC1 and cDC2 subsets of cDCs are activated and migrate equally well in response to TLR3 stimulation in a cell extrinsic and TNFα dependent manner, but that cDC1 show a unique requirement for type I interferon signaling. Our findings reveal common and differing pathways regulating DC subset migration, offering important insights for the design of DC-based vaccination and therapy approaches.

## Introduction

Dendritic cells (DCs) are the major antigen-presenting cells in the body, which, upon migration to secondary lymphoid organs, initiate and shape naïve T cell responses to peripherally acquired antigen. DCs are divided into two major subsets referred to as cDC1 and cDC2 (Guilliams et al., 2014). In the intestine, migratory cDC1 are defined as XCR1^+^CD103^+^CD11b^−^, while cDC2 can be divided into a major XCR1^−^CD103^+^CD11b^+^ and a minor XCR1^−^CD103^−^CD11b^+^ subset. Although both subsets present mucosa-derived antigen in the draining lymph nodes (LNs), cDC1 and cDC2 differ in their capacity to induce specific immune responses (Eisenbarth, 2018). While cDC1 are generally implicated in viral defense and cross presentation of exogenous antigens to MHCI-restricted CD8^+^ T cells and MHCII-restricted CD4^+^ T_H_1 cells (Hildner et al., 2008), cDC2 are highly effective at inducing T_H_17 and T_H_2 responses (Persson et al., 2013; Schlitzer et al., 2013; Williams et al., 2013). Specific targeting of DC subsets is thus of high relevance for DC-based strategies for vaccination and therapeutic approaches against different types of antigen.

Antigen-targeting to specific DC subsets using antibody-mediated delivery to differentially expressed surface receptors can indeed shape the resulting type of immunity (Dudziak et al., 2007). One family of molecules expressed differentially by DC subsets is toll-like receptors (TLRs)(Denning et al., 2011; Edwards et al., 2003), suggesting that differential engagement of DC subsets could also be achieved by using adjuvants specifically activating one subset but not the other. In support of this idea, the induction of fully functional cytotoxic CD8^+^ T lymphocytes depends on simultaneous uptake of antigen together with cell-intrinsic stimulation of pattern recognition receptors expressed by the presenting DC (cis-activation)(Desch et al., 2014). TLR3 is an endosomal receptor that recognizes double-stranded RNA (dsRNA), a molecular pattern associated with viral infections (Alexopoulou et al., 2001; Matsumoto et al., 2002). As several studies have demonstrated that TLR3 is preferentially expressed in cDC1 (Davey et al., 2010; Edwards et al., 2003; Jelinek et al., 2011; Luber et al., 2010) and promotes cross-presentation of antigen with high efficiency (Mandraju et al., 2014; Rizzo et al., 2016; Schulz et al., 2005), targeting TLR3 is a promising strategy in cancer-immunotherapy and vaccination against viruses. A hallmark of DCs is to migrate to the draining LNs to present peripherally acquired antigen. In response to the TLR7-stimulating agent R848, plasmacytoid (pDC)-derived TNFα drives cDC migration from the small intestinal lamina propria (SI LP) to the mesenteric LNs (mLNs), while type I interferon (IFN) regulates DC activation (Yrlid et al., 2006). Subset-specific requirements were not assessed. Most TLR3 driven transcriptional changes in splenic DCs after stimulation with the double-stranded (ds)RNA mimic polyinosinic:polycytidylic acid (poly(I:C)) result from secondary effects through the type I IFN receptor on cDCs (Pantel et al., 2014). This suggests that migration in response to poly(I:C) may also depend on type I IFN signaling. Here we have analyzed in detail the major cellular and molecular players involved in the activation and migration of intestinal cDC subsets in response to poly(I:C) *in vivo* and provide novel insights regarding cis- and trans-regulation of these processes.

## Results and Discussion

### Poly(I:C)-induced intestinal DC migration depends on TLR3 signaling

We first set out to analyze in detail the expression of TLR3 by immune cells of the spleen, mLNs and small intestine lamina propria (SI-LP) and confirmed that only cDC1 DCs expressed high amounts of TLR3 in all organs (Fig. 1A). While macrophages also expressed low levels of TLR3, cDC2 were almost entirely negative and B and T cells showed no expression (Fig. 1A and Supplemental Fig. 1A). Importantly, stimulation with poly(I:C) did not change TLR3 expression across subsets (Fig. 1A). These results are consistent with *in vitro* data on the differential abilities of the cDC subsets to poly(I:C) stimulation (Jelinek et al., 2011) and therefore we hypothesized that poly(I:C) would drive migration of cDC1 preferentially *in vivo.* To test this, we quantified CD103^+^ cDC1 and cDC2 in the mLN after intraperitoneal injection of poly(I:C), based on the knowledge that CD103 expression by cDC in the mLN defines those which are derived from CCR7 dependent migration from the intestinal mucosa (Hagerbrand et al., 2015; Johansson-Lindbom et al., 2005). Consistent with this idea, the numbers of both migratory cDC1 and cDC2 increased after administration of poly(I:C), peaking at 12 hours post-injection and returning to steady state levels after 24h (Fig. 1B). Interestingly, cDC2 migrated almost as efficiently as cDC1, with only a small disadvantage being seen at early time points.

**Fig.1:**
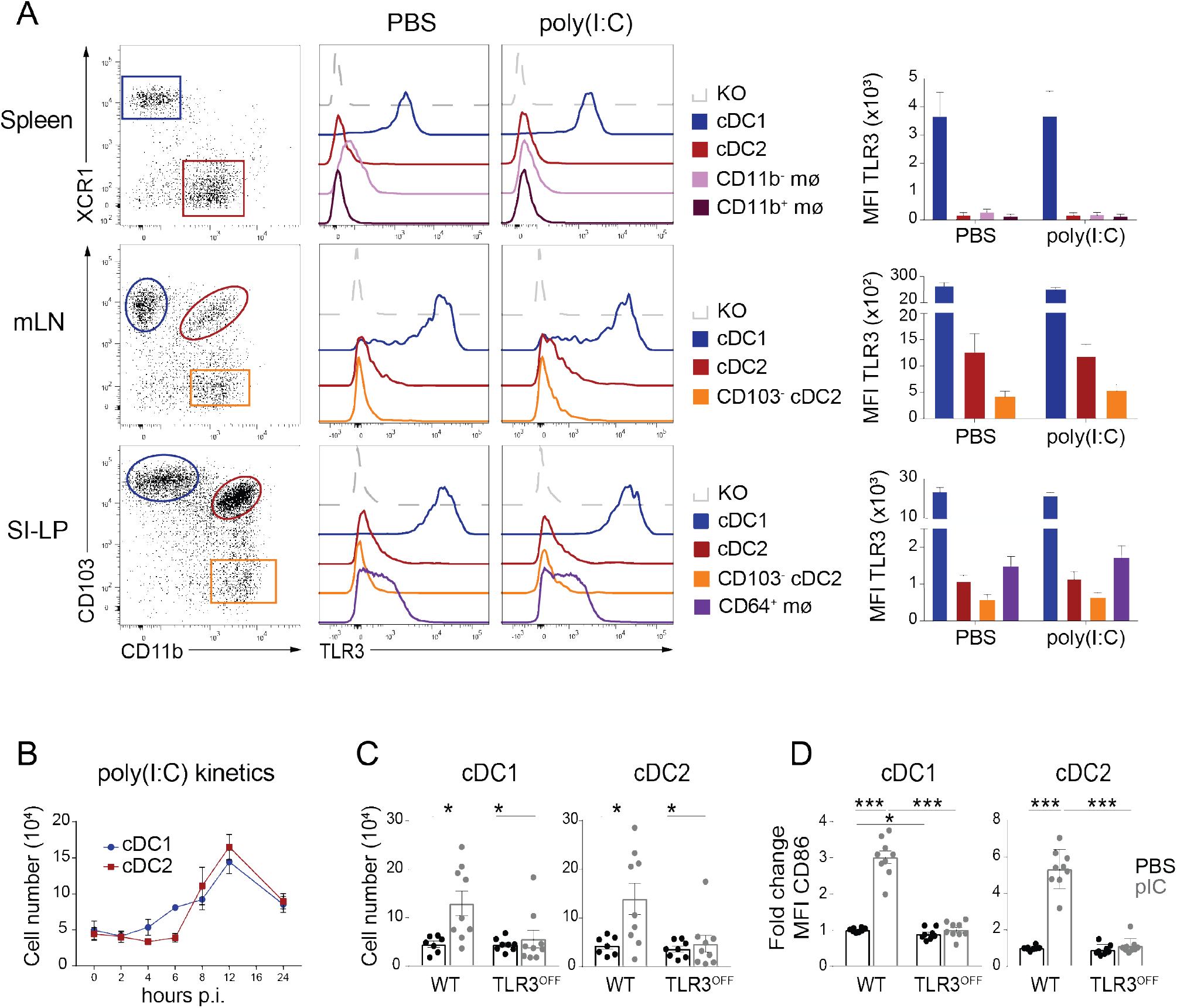
TLR3 expression by mononuclear phagocytes and migration of cDCs in response to poly(I:C). A. Left: representative flow cytometry plots of spleen, mLN and SI-LP DC subsets and macrophages in C57BL/6 mice. All populations were gated on live, lineage (CD3, CD19, NK1.1) negative, single cells. DC in spleen and mLN were further pre-gated as CD11c^+^MHCII^+^ cells and in SI-LP as CD11c^+^MHCII^+^CD64^−^ cells. Macrophages in spleen were further pre-gated as CD11c^int^ and CD11b^+^ or CD11b^−^, while macrophages in SI-LP were further gated as CD11c^+^CD64^+^ cells. Histograms: Intracellular TLR3 staining of the indicated DC and macrophage populations 12 hours after i.p injection of PBS or 100μg poly(I:C) into wild type mice and by bulk DC in resting TLR3^OFF^ mice (KO). Right: Quantification of TLR3 expression by DC subsets and macrophages in C57BL/6 mice 12h after i.p. injection of PBS or poly(I:C). Data shown are means ± 1 sem pooled from two independent experiments with 3 mice per group. B. Kinetics of intestinal cDC1 and cDC2 migration after i.p. injection of 100μg poly(I:C). Data shown are mean numbers of cells ± 1 sem pooled from two to four independent experiments with 2-3 mice per group. Differences between cDC1 and cDC2 are not significant. C. Total numbers of cDC1 and cDC2 in the mLNs of WT and TLR3^OFF^ mice 12h after i.p. injection of PBS or 100μg poly(I:C). Data shown are mean numbers of cells ± 1 sem pooled from three independent experiments with 3 mice per group. Two-way ANOVA, *p<0.05. D. Activation of cDC subsets in mLNs of WT and TLR3^OFF^ mice by poly(I:C). Results shown are fold change in CD86 expression 12h after injection of 100μg poly(I:C) as assessed by MFI normalized to FMO and relative to expression by DCs in untreated WT. Data shown are means ± 1 sem pooled from three independent experiments with 3 mice per group. Two-way ANOVA, *p<0.05, ***p<0.0005.

Although retinoic acid-inducible gene 1 (RIG-I)-like helicases that signal through mitochondrial antiviral-signaling protein (MAVS) can also sense poly(I:C) (Jensen and Thomsen, 2012), DC migration of both subsets was completely abrogated in TLR3-deficient mice (Fig. 1C) and in mice deficient for the TLR3 adapter TRIF (TIR-domain-containing adapter-inducing interferon-β) (Supplemental Fig. 1B). As DC migration and activation, both crucial events for the induction of immunity, can be regulated independently (Jones et al., 2016; Yrlid et al., 2006), we also measured the expression of the costimulatory molecule CD86 as a surrogate marker for DC activation. Again, activation of both DC subsets was also entirely depended on TLR3 and TRIF expression (Fig. 1D and Supplemental Fig. 1C), showing that poly(I:C) induces migration and activation of both cDC1 and cDC2 in a strictly TLR3-dependent manner. Our findings are in accordance with previously published data showing that *in vitro* activation with poly(I:C) is abrogated in bone-marrow (BM)-derived DCs from TLR3-deficient mice (Jelinek et al., 2011). However as cDC2 themselves express virtually no TLR3, our data indicate that TLR3 stimulation might act in both cell-intrinsic and extrinsic manners on cDCs *in vivo.*

### Cell-intrinsic TLR3-sensing is dispensable for DC migration

Non-hematopoietic cells express TLR3 and support immune cell survival, maturation and function. For example TLR3 expression in intestinal epithelial cells is required for optimal clearance of rotavirus (Pott et al., 2012) and epithelial cells have previously been implicated in driving DC migration to the draining LNs during viral infection (Ye et al., 2019). To determine whether TLR3-dependent sensing in non-hematopoietic cells could induce intestinal DC migration in response to poly(I:C), we reconstituted irradiated wild-type mice with TLR3-deficient BM and treated the mice with poly(I:C). The results showed that while DCs migrated well in response to poly(I:C) in WT recipients of WT BM, there were no significant increases in mLN DC numbers in recipients of TLR3-deficient BM after administration (Fig. 2A). Thus TLR3-expression within the hematopoietic compartment is required to drive efficient DC migration in response to poly(I:C).

**Fig. 2:**
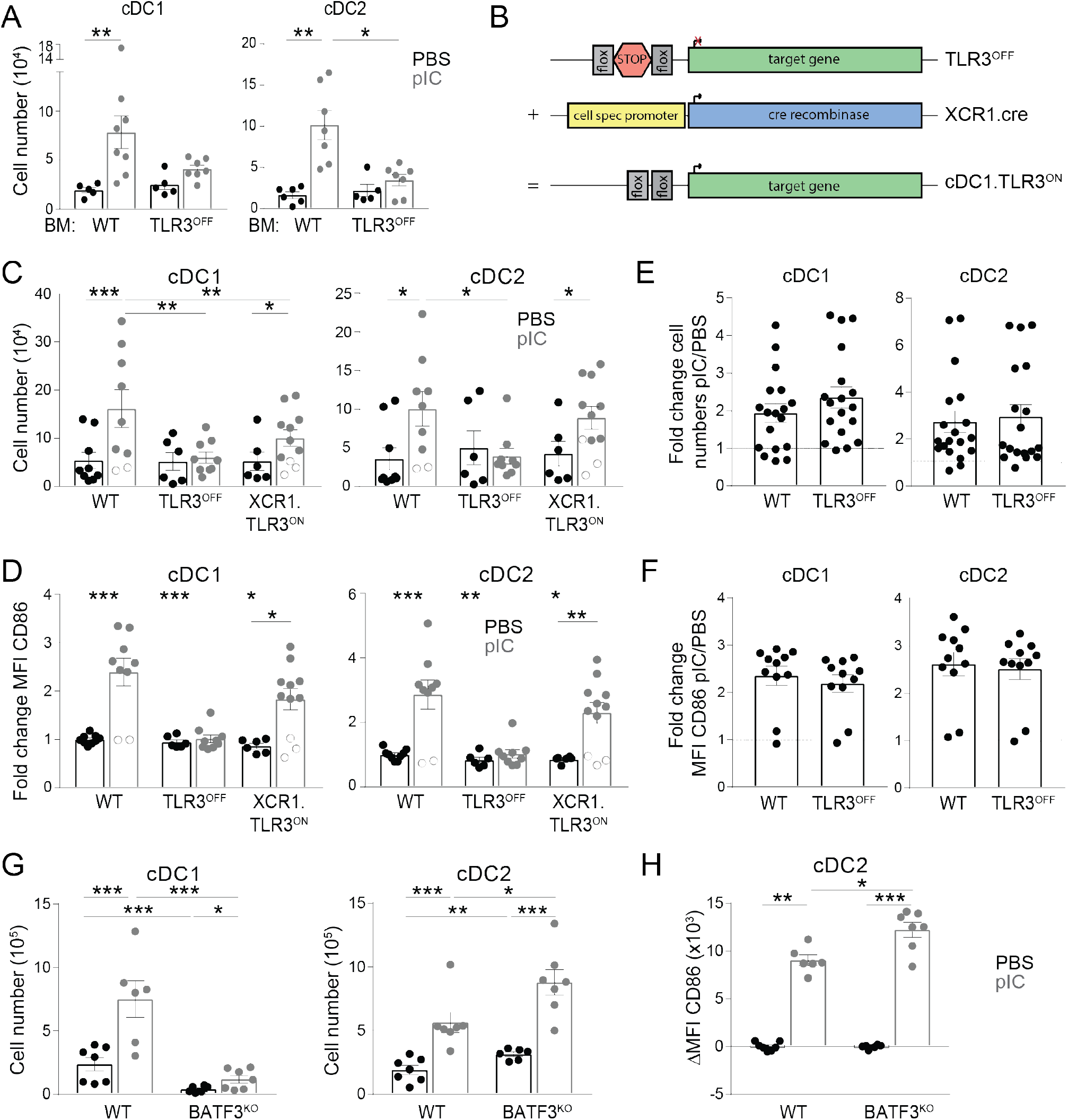
Cellular requirements for TLR3 mediated DC migration in response to poly(I:C) A. Total numbers of cDC1 and cDC2 in the mLNs of WT recipients reconstituted for 8 weeks with either WT or TLR3^OFF^ BM 12h after i.p. injection of PBS or 100μg poly(I:C). Data shown are mean numbers of cells ± 1 sem from one experiment with 5-8 mice per group. Mann Whitney U test,*p<0.05, **p<0.005. B. Schematic diagram of generation of cell specific TLR3^ON^ mice in which TLR3^OFF^ was created using a floxed STOP codon and TLR3 then re-expressed using cell specific cre promoters to delete the STOP codon. C. Total numbers of cDC1 and cDC2 in the mLNs of WT, TLR3^OFF^ and XCR1.TLR3^ON^ mice 12h after i.p. injection of PBS or 100μg poly(I:C). Data shown are mean numbers of cells ± 1 sem pooled from three independent experiments with 3-4 mice per group. Two-way ANOVA, *p<0.05, **p<0.005, ***p<0.0005. Open circles were used to mark those poly(I:C) injected mice that also did not show upregulation of CD86 (panel D); these were not excluded from statistics. D. Activation of cDC subsets in mLNs of WT, TLR3^OFF^ and XCR1.TLR3^ON^ mice by poly(I:C). Results shown are fold change in CD86 expression 12h after i.p. injection of 100μg poly(I:C) as assessed by MFI normalized to FMO and relative to expression by DCs in untreated WT. Data shown are means ± 1 sem pooled from three independent experiments with 3-4 mice per group. Two-way ANOVA, *p<0.05, **p<0.005, ***p<0.0005. E. Fold change of total number of cDC1 and cDC2 in the mLNs 12h after i.p. injection of 100μg poly(I:C) versus PBS derived from the indicated BM in 50:50 WT:TLR3^OFF^ mixed BM chimeras. Data shown are means ± 1 sem pooled from two independent experiments with 7 mice per group. Two-way ANOVA, not significant. F. Activation of cDC subsets in mLNs of 50:50 WT:TLR3^OFF^ mixed BM chimeras by poly(I:C). Results shown are fold change in CD86 expression 12h after i.p. injection of 100μg poly(I:C) versus PBS and relative to expression by DCs in untreated WT. Data shown are means ± 1 sem pooled from two independent experiments with 3-7 mice per group. Two-way ANOVA, not significant G. Total number of cDC1 and cDC2 cells in BATF3^KO^ mice 12 after i.p. injection of PBS or 100 μg poly(I:C). Data shown are mean numbers of cells ± 1 sem pooled from two representative experiment out of 3 with 2-4 mice per group. Two-way ANOVA, *p<0.05, **p<0.005, ***p<0.0005. H. Activation of cDC2 subset in mLNs of BATF3^KO^ mice by poly(I:C). Results are shown as mean fluorescent intensity of CD86 12h after i.p. injection of PBS or 100μg poly(I:C) in WT and BATF3^KO^ mice. Two-way ANOVA, *p<0.05, **p<0.005, ***p<0.0005.

As cDC1 uniformly expressed TLR3, we explored the role of this subset in sensing poly(I:C) for driving DC migration directly, by generating a mouse model that allows for cell-specific re-expression of TLR3 in a TLR3 KO background. To this end, a floxed transcriptional termination cassette was inserted into the coding sequence of the TLR3 gene (TLR3^OFF^), abolishing TLR3 expression. Expression of TLR3 by cDC1s could then be restored in cDC1.TLR3^ON^ mice in which the TLR3 stop codon was deleted using XCR1-driven cre recombinase (XCR1.cre (Janela et al., 2019)) (Fig. 2B and Supplemental Fig. 2A). Poly(I:C)-induced DC migration and activation of both cDC1 and cDC2 occurred in cDC1.TLR3^ON^ mice, but to a lesser extent compared to WT mice (Fig. 2C,D). As expected, DC migration was absent in TLR3^OFF^ mice (Fig. 2C,D). These findings suggest that while cDC1-restricted TLR3 expression can drive poly(I:C) induced DC migration, other TLR3-expressing cells contribute to optimal DC migration in response to poly(I:C). Of note, careful analysis of XCR1-driven re-expression of TLR3 revealed that re-expression of TLR3 also occurred in ~20% of CD64^+^CD11b^+^XCR1^−^ macrophages in the intestine, but not in spleen macrophages (Supplemental Fig. 2A). This phenomenon is not specific for the TLR3 locus, as XCR1.cre could also drive YFP expression by some intestinal macrophages when crossed to ROSA-STOP-YFP (data not shown). We therefore examined whether off-target re-expression of TLR3 by intestinal macrophages might account for the restored DC migration in cDC1.TLR3^ON^ mice. However, migration of both cDC1 and cDC2 was entirely normal after administration of poly(I:C) to CCR2-deficient mice that lack most of the monocyte-derived intestinal macrophages (Bain et al 2014) (Supplemental Fig. 2B). Finally, we could not detect any migration or activation of either DC subset if TLR3 expression was restricted to intestinal epithelial cells of TLR3^OFF^ mice using villin-cre (villin.TLR3^ON^, Supplemental Fig. 2C). Together, these data suggest that cDC1-specific TLR3 expression can drive DC migration in response to poly(I:C), although a contributory role for a residual population of CCR2-independent, TLR3 expressing intestinal macrophages in cDC1.TLR3^ON^ mice is likely.

The fact that expression of TLR3 restricted to XCR1-expressing cDC1 can drive the migration and activation of cDC2 indicates a cell extrinsic effect of poly(I:C) on cDC2. To test whether cell extrinsic effects on cDC1 are equally sufficient, we generated mixed-BM chimeras in which WT recipients on a CD45.1/.2 congenic background were reconstituted with a 50:50 mix of CD45.1^+^ WT and CD45.2^+^ TLR3-deficient BM. Under these conditions, administration of poly(I:C) induced the activation and migration of TLR3-deficient cDC1 and cDC2 to the same extent as their WT counterparts in the same hosts (Fig. 2E,F), indicating that both cDC1 and cDC2 can respond to TLR3 stimulation in a cell extrinsic manner. This is presumably driven by the TLR3-competent bone marrow derived cells of WT origin present in the mixed chimeras. Interestingly, cDC1 themselves do not appear to play an essential role in this process, as complete deficiency of cDC1 DCs in BATF3^KO^ mice (Edelson et al., 2010) did not abrogate the activation and migration of cDC2 in response to poly(I:C), showing that hematopoietic cells other than cDC1 can also contribute (Figure 2G,H). Macrophages are a potential candidate for this role, as they express and respond to TLR3-stimulation (Zhou et al., 2010) and we attempted to explore their involvement by generating macrophage-specific TLR3^ON^ mice using LysM.cre (McCubbrey et al., 2017) to delete the TLR3 stop codon in TLR3^OFF^ mice. However this approach was unsuccessful, as TLR3 was re-expressed by ~50% of cDC1 of LysM.TLR3^ON^ mice and thus the role of macrophages in responding to TLR3 *in vivo* requires further investigation (Supplemental Fig. 2D).

Taken together, these results show that the hematopoietic compartment is responsible for TLR3-dependent migration and activation of DCs, but that these processes can occur in a cell-extrinsic manner, with cDC1-derived signals not being essential, despite the high levels of TLR3 expression by these cells.

### DC migration in response to poly(I:C) is independent of MyD88, but requires TNF receptor signaling

The cell-extrinsic effect of TLR3 on DC migration in response to poly(I:C) suggests that inflammatory mediators produced following TLR3 signaling on TLR3^+^ target cells might play a key role in this process. We therefore measured the expression of cytokines that have been implicated in DC activation and migration by qPCR analysis of whole SI tissue samples at different times after administration of poly(I:C). This showed increased levels of mRNA for TNF-α, IL-1β, IFN-α, and IFN-β after 2 and 4 hours after poly(I:C) injection (Figure 3A).

**Fig.3:**
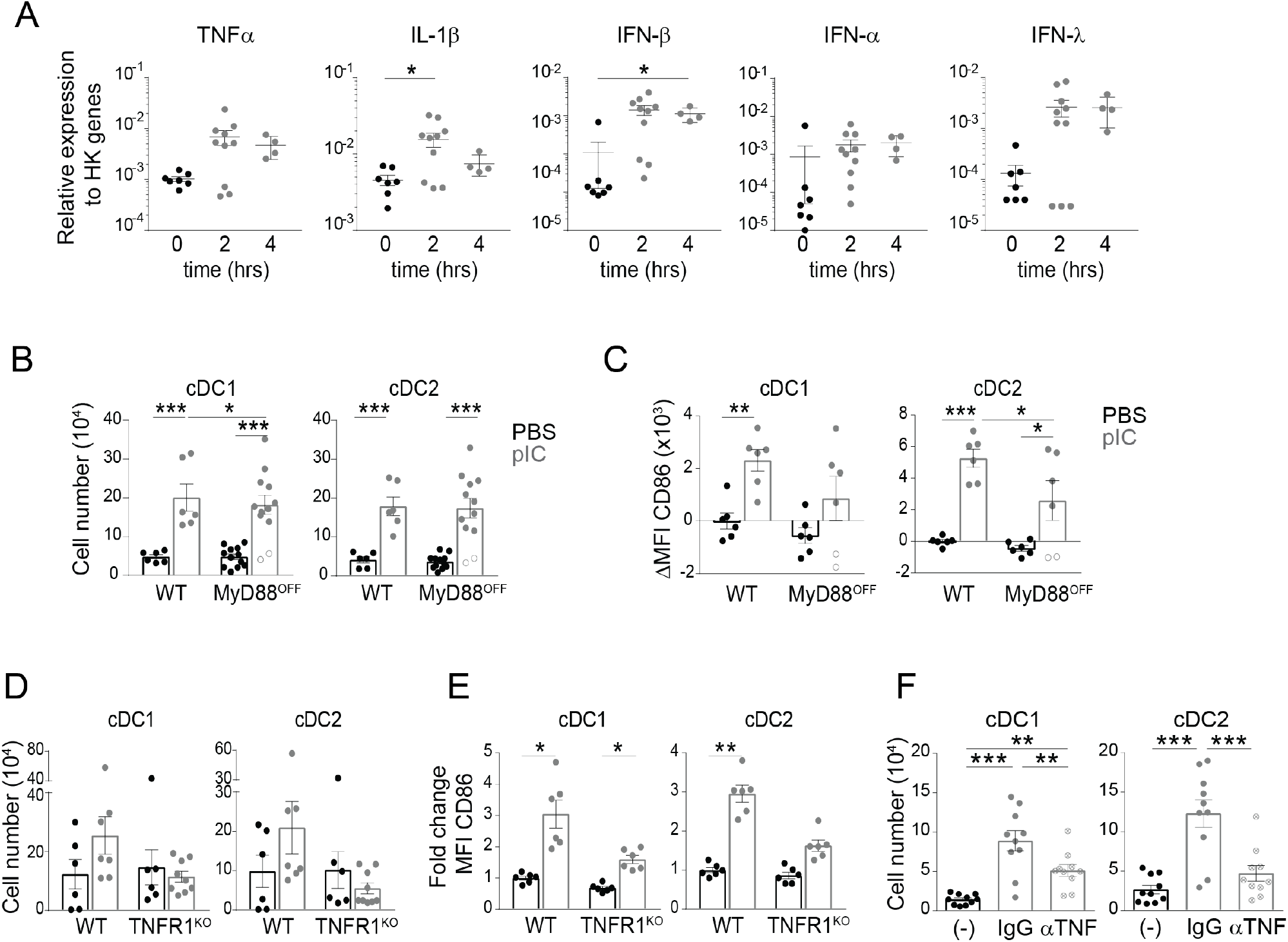
Role of cytokines and MyD88 in response of DC to poly(I:C) A. Expression of cytokine mRNA in total SI LP of WT C57BL/6 mice at indicated times after i.p. injection of 100μg poly (I:C). Each point represents mean of qPCR triplicates for every gene as assessed by RT-qPCR and measured relative to mean of three housekeeping genes (Reep5, β-actin, GAPDH). Data shown are means ± 1 sem pooled from three independent experiments with 2-4 mice per group. Two-way ANOVA, *p<0.05. B. Total number of cDC1 and cDC2 in the mLNs of WT and MyD88^OFF^ 12h after i.p. injection of PBS or 100μg poly(I:C). Data shown are mean numbers of cells ± 1 sem pooled from four independent experiments with 3-4 mice per group (only two including WT). Two-way ANOVA, *p<0.05, ***p<0.0005. Open circles were used to mark those poly(I:C) injected mice that also did not show upregulation of CD86 (panel C); these were not excluded from statistics. C. Activation of cDC subsets in mLNs of WT and MyD88^OFF^ mice by poly(I:C). Results shown are delta MFI of CD86 expression 12h after i.p. injection of PBS or 100μg poly(I:C) over the mean of all untreated WT CD86 MFI values. Data shown are means ± 1 sem pooled from two independent experiments with 3-4 mice per group. Two-way ANOVA, *p<0.05, **p<0.005, ***p<0.0005. D. Total number of cDC1 and cDC2 in the mLNs of TNFR1^KO^ mice 12h after i.p. injection of PBS or 100μg poly(I:C). Data shown are mean numbers of cells ± 1 sem pooled from three independent experiments with 1-3 mice per group. Two-way ANOVA, not significant. E. Activation of cDC subsets in mLNs of WT and TNFR1^KO^ mice by poly(I:C). Results shown are fold change in CD86 expression 12h after injection of 100μg poly(I:C) as assessed by MFI normalized to FMO and relative to expression by DCs in untreated WT. Data shown are means ± 1 sem pooled from three independent experiments with 1-3 mice per group. Two-way ANOVA, *p<0.05, **p<0.005. F. Total number of cDC1 and cDC2 in the mLNs of C57BL/6 mice pre-treated with TNF-α antibody-blocking and 12h after i.p. injection of 100μg poly(I:C). Control mice were treated with the isotype antibody (IgG). Data shown are mean numbers of cells ± 1 sem pooled from three independent experiments with 2-4 mice per group. Two-way ANOVA, **p<0.005, ***p<0.0005

Steady state migration of intestinal DCs depends on MyD88 signaling through NFκB (Baratin et al., 2015; Hagerbrand et al., 2015) and although TLR3 signaling itself does not require MyD88, the IL1 receptor signals through MyD88 (Dinarello, 2009). However, the activation and migration of both cDC1 and cDC2 occurred normally in poly(I:C) treated MyD88^KO^ mice (Figure 3B, C). Although TNF receptor 1 (TNFR1) signaling is not important for intestinal DC migration in the steady state, it is required for DCs to migrate in response to R848 (Hagerbrand et al., 2015). TNFR1 signaling is important for the induction of the CD8 T cell response towards mouse hepatitis virus, and expression on DCs alone is sufficient to confer protection (Ding et al., 2011). As we found TNFα to be upregulated in the intestine after injection with poly(I:C), we examined its role in poly(I:C) induced DC migration. Indeed, there were no significant increases in migration of either cDC1 or cDC2 in response to poly(I:C) in either TNFR1^KO^ mice or mice treated *in vivo* with a blocking anti-TNFR1 antibody, while there was minimal DC activation in poly(I:C) treated TNFR1^KO^ mice (Figures 3D-F). These results are consistent with previous studies on skin DCs (Suto et al., 2014) and show that TNFR1 signaling is a crucial secondary signal that mediates the extrinsic response of DCs to TLR3-sensing *in vivo.*

### DC subsets differ in type I IFN signaling requirements for migration and activation in response to poly(I:C)

In addition to elevated expression of TNF-α and IL-1β, type I IFNs were significantly upregulated in the intestine after poly(I:C) injection (Figure 3A). Previous studies have shown a prominent role for type I IFN in the activation and maturation of splenic DCs in response to poly (I:C), acting via the type I IFN receptor on DCs (Pantel et al., 2014). Conversely, the TNFα dependent migration of intestinal DCs in response to R848 does not require type 1 IFNR signaling but it is rather needed for their activation (Yrlid et al., 2006). We therefore tested directly the role of type I IFN in the activation and migration of intestinal DCs in response to poly(I:C).

Type I IFN receptor-deficient mice (IFNAR^KO^) showed defective migration of both cDC1 and cDC2 in response to poly(I:C) (Figure 4A) and similar results were found in mice lacking IFNAR in all CD11c-expressing cells, although cDC2 showed some residual migration upon deletion of IFNAR in these mice (Figure 4B). The activation of both DC subsets as assessed by CD86 expression was also greatly diminished in CD11c.IFNAR^KO^ mice (Figure 4C). Conversely, while cDC1-specific deletion of the IFNAR in XCR1.IFNAR1^KO^ mice abrogated the poly(I:C) induced migration and activation of cDC1, this had no effect on either parameter in cDC2 (Figures 4B, C). Deletion of IFNAR in cDC2 in huCD207.IFNAR1^KO^ mice however had no effect on the migration or activation of either DC subset, apart from a small decrease in CD86 upregulation by cDC2 (Figures 4B, C). These data suggest that type I IFN signaling in cDC2 is not required for their migration to the mLNs in response to poly(I:C).

**Fig.4:**
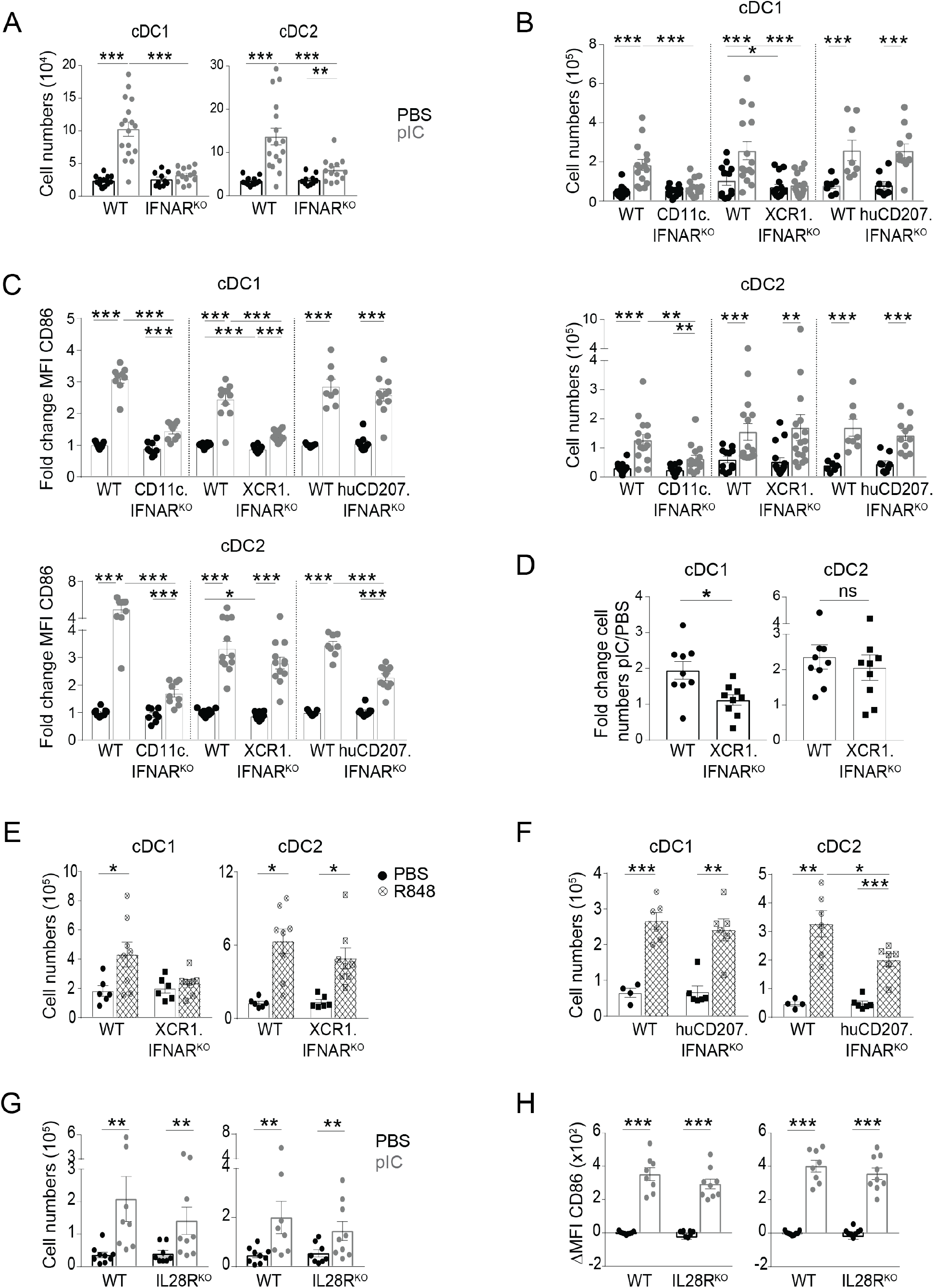
Type I IFN signaling in migration and activation of DC subsets. A. Total number of cDC1 and cDC2 cells in the mLNs of WT and IFNAR^KO^ mice 12h after i.p. injections of PBS or 100μg poly(I:C). Data shown are mean numbers of cells ± 1 sem pooled from four independent experiments with 3-5 mice per group. Two-way ANOVA, **p<0.005, ***p<0.0005. B. Total number of cDC1 (top) and cDC2 (bottom) cells in the mLNs of WT, CD11c.IFNARKO^KO^, XCR1.IFNAR^KO^ and huCD207.IFNAR^KO^ mice 12h after i.p. injection of PBS or 100μg poly(I:C). Data shown are mean numbers of cells ± 1 sem pooled from five independent experiments with 2-3 mice per group for WT vs CD11c.IFNAR^KO^; five independent experiments with 3-5 mice per group for XCR1.IFNAR^KO^, and three independent experiments 3-5 mice per group for huCD207.IFNAR^KO^. Two-way ANOVA within littermates, *p<0.05, **p<0.005, ***p<0.0005. C. Activation of cDC1 (top) and cDC2 (bottom) in the mLNs of WT, CD11c.IFNAR^KO^, XCR1.IFNAR^KO^ and huCD207.IFNAR^KO^ mice by poly(I:C). Results shown are fold change in CD86 expression 12h after i.p. injection of 100μg poly(I:C) as assessed by MFI normalized to FMO and relative to expression by DCs in untreated WT. Data shown are means ± 1 sem pooled from three out of five independent experiments with 2-3 mice per group for WT vs CD11c.IFNAR^KO^; four out of five independent experiments with 3-5 mice per group for XCR1.IFNAR^KO^, and three independent experiments 3-5 mice per group for huCD207.IFNAR^KO^. Two-way ANOVA within littermates, *p<0.05, ***p<0.0005. D. Fold change of total number of cDC1 and cDC2 in the mLNs 12h after i.p. injection of 100μg poly(I:C) versus PBS derived from the indicated BM in 50:50 WT:XCR1.IFNAR^KO^ mixed BM chimeras. Data shown are means ± 1 sem pooled from two independent experiments with 3-9 mice per group. Two-way ANOVA, *p<0.05. E. Total number of cDC1 and cDC2 in the mLNs of XCR1.IFNAR^KO^ mice 12h after oral gavage of PBS or 20μg R848. Data shown are mean numbers of cells ± 1 sem pooled from two independent experiments with 3-5 mice per group. Two-way ANOVA, *p<0.05. F. Total number of cDC1 and cDC2 in the mLNs of huCD207.IFNAR^KO^ mice 12h after oral gavage of PBS or 20μg R848. Data shown are mean numbers of cells ± 1 sem pooled from two independent experiments with 3-5 mice per group. Two-way ANOVA, *p<0.05, **p<0.005, ***p<0.0005. G. Total number of cDC1 and cDC2 in the mLNs of IL28R^KO^ mice 12h after i.p. injection of PBS or 100μg poly(I:C). Data shown are mean numbers of cells ± 1 sem pooled from three independent experiments with 3 mice per group. Two-way ANOVA, **p<0.005. H. Activation of cDC1 and cDC2 in the mLNs of IL28R^KO^ mice by poly(I:C). Results shown are delta MFI of CD86 expression 12h after i.p. injection of PBS or 100μg poly(I:C) over the mean of all untreated WT CD86 MFI values. Data shown are means ± 1 sem pooled from three independent experiments with 3 mice per group. Two-way ANOVA, ***p<0.0005.

Since we observed that TLR3-signaling was not required cell-intrinsically (Figure 2E), we next checked whether this was also the case for type IFN signaling. Mixed BM chimeras using a 50:50 combination of WT and XCR1.IFNAR1^KO^ BM showed that the requirement for type I IFN signaling in cDC1 migration was cell intrinsic (Figure 4D). Poly(I:C) induced migration of cDC2 remained intact in these chimeras (Figure 4D). A similar defect in cDC1 migration was seen in XCR1.IFNAR1^KO^ mice treated orally with R848, the ligand for TLR7 expressed mostly by cDC2, indicating a need for type I IFN signaling in cDC1 regardless of whether stimulation occurred in a direct or indirect manner (Figure 4E). cDC2 migration was also induced by R848 in huCD207.IFNAR1^KO^ mice lacking type I IFN signaling specifically on cDC2, although this was somewhat reduced in comparison with that in WT mice (Fig. 4F). Thus, type I IFN signaling plays a global role in the migration of cDC1 in response to TLR stimulation, but has little or any effect on cDC2 migration in response to either TLR3 or TLR7 stimulation.

As well as type I IFN, poly(I:C) also induces cDC1 to express IFN-λ in a manner that requires IFNAR on splenic DCs (Lauterbach et al., 2010). IFN-λ is a type III IFN that drives thymic stromal lymphopoietin expression by M cells in response to nasal vaccination with an influenza vaccine, which in turn drives cDC1 migration from the respiratory tract to the mediastinal lymph nodes (Ye et al., 2019). We therefore tested whether IFN-λ was required for the poly(I:C) induced migration of DCs, using mice deficient for IL28R, the receptor for IFN-λ. However the activation and migration of both cDC1 and cDC2 in response to poly(I:C) were normal in these animals (Figure 4G,H).

Our data reveal a previously unappreciated differential role for type I IFN in cDC migration from the intestine to the mLNs. While IFNAR signaling drives maturation of both major subsets of migratory DCs, only cDC1 critically depend on direct type I IFN signals for migration in response to poly(I:C). Thus, the mechanisms inducing DC migration may be subset specific. Although the migration and upregulation of CD86 in both cDC1 and cDC2 were entirely TLR3-dependent, this occurred in a cell-extrinsic manner and the cells responding directly to TLR3 remain to be identified. Consistent with previous reports in other models ((Longhi et al., 2009; Pantel et al., 2014; Yrlid et al., 2006); we found that TNFα and type I IFN signaling played important roles as secondary mediators in TLR3-mediated intestinal DC migration. Interestingly, the ability of DCs to induce proliferation by naïve CD4^+^ T cells also does not require cell intrinsic expression of pattern recognition receptors by DCs, whereas the functional polarization of T cells depends on direct sensing of the pathogen-associated molecular pattern by the presenting DC (Desch et al., 2014; Spörri and Reis e Sousa, 2005). Our data showing that the migration and upregulation of costimulatory molecules by DCs in response to TLR stimuli *in vivo* can result from trans-activation of DCs therefore suggest that differences in migration capacities may not be linked to the polarization of T cell responses. However it is important to note that not all TLR ligands can induce migration of both DC subsets, as signaling through TLR5, that is only expressed on cDC2, does not induce potent cDC1 migration *in vivo* (Flores-Langarica et al., 2017). Taken together, these findings indicate that TLR based adjuvants and targeting need to be examined individually for their impact on specific DC subsets if their effects on the immune system are to be understood in depth. Future research aiming at better understanding the role of potently migrating and activating trans-activated DC subsets, as well as the signals responsible, will be critical for the design of selective immune interventions in vaccination and therapy.

## Acknowledgements

We thank Prof. Dan Kaplan for sharing huCD207-cre mice and Prof. Ulrich Kalinke for sharing IFNAR^flox^ mice. We thank the Lahl, Agace, Bekiaris and Johansson-Lindbom laboratories for many fruitful discussions, Matthias Schiemann and Immanuel Andrä for flow cytometry support, and Allan Mowat for editing of the manuscript. KL was supported by a Vetenskaprådet Young Investigator Award, the Ragnar Söderberg Foundation Fellowship in Medicine, a Lundbeck Foundation Research Fellowship, the Åke Wiberg Foundation, the Carl Trygger Foundation, and the Crafoord Foundation. JH was supported by a MARIE CURIE Fellowship as part of the EU-funded project HC Ørsted Postdoc. AGL received travel support from the Niels Bohr Foundation. VB and WA were supported by a project grant from Vetenskaprådet. The flow cytometer at Lund University was purchased with funding from the Lundberg Foundation.

## Author contributions

AGL, VB, WA, and KL conceived and designed the project; AGL, VB, JH, MO’K, WA and KL designed experiments; AGL, VB, KM-L, JH, KGM, JN, IU, KK, and KL performed experiments and analysed the data; BM and BH provided essential tools; AGL and KL wrote the manuscript; WA and KL supervised the study and acquired funding; all authors reviewed the manuscript.

## Experimental procedures

### Mice

All animal animals were house under specific pathogen-free conditions at the Danish Technical University (Denmark), Lund University (Sweden) or Monash University (Australia). The experiments were performed under the appropriate national licenses and guidelines for animal care. Both male and female mice were used between 8 and 16 weeks of age as no obvious age differences were detected. CD11c.cre mice (B6.Cg-Tg(Itgax-cre)1-1Reiz/J (Caton et al., 2007)) allow floxed gene deletion in CD11c-expressing cells, huCD207.cre mice drive floxed gene deletion in Langerhans cells and intestinal cDC2 (Welty et al., 2013), XCR1.cre mice permit to specifically delete floxed genes in cDC1 (Janela et al., 2019), villin.cre (B6.Cg-Tg(Vil-cre)997Gum/J) mice excise floxed genes in intestinal epithelial cells (Madison et al., 2002) and Rosa26-STOP-YFP mice allow tracking of cre specificity (B6.129X1-Gt(ROSA)26Sor^tm1(EYFP)Cos^/J). We used “switch-on” mutants carrying a floxed stop cassette in the endogenous locus prior to the gene of interest, allowing for re-expression of the targeted gene in the presence of cre for MyD88 (Gais et al., 2012), TLR3 and TRIF (unpublished, manuscript in preparation) (both generated at TU Munich, Germany). IFNAR floxed mice were obtained from U. Kalinke (Kamphuis et al., 2006). BATF3^KO^ (B6.129S(C)-Batf3^tm1Kmm^/J) were maintained at DTU, TNFR1^KO^ (crossed out from TNFR1/2^KO^ (Peschon et al., 1998)), IFNAR^KO^ (Cucak et al., 2009) and CCR2^KO^ (B6.129S4-Ccr2^tm1Ifc^/J) at Lund University and IL28RA^KO^ (Ank et al., 2008) (kindly provided by Sean Doyle, Zymogenetics/BMS) at Monash University. All mice were on the C57Bl/6J background (B6.SJL-Ptprc^a^Pepc^b^/BoyJ for CD45.1 bone marrow donors) and littermates were used as controls.

### In vivo treatments

Mice were injected with PBS or 100μg pIC (Sigma-Aldrich) in PBS intraperitoneally (i.p.) and mLNs and small intestinal lamina propria (SI LP) were collected 12-14 h later if not indicated otherwise. For TLR7 stimulation, 20μg of R848 (Invivogen) in PBS was given orally. αTNFα (XT3.11, BioXcell) was blocked with 0.5mg on day −1 and 0.5mg at the time of stimulation.

### Cell isolation

Isolation of mLN and splenic DCs was performed by digesting the tissue with collagenase IV (0.5 mg/mL, Sigma-Aldrich) and DNase I (12.5 μg/mL) diluted in R10 media (RPMI 1640 + 10% FCS) for 40 min at room temperature. Remaining tissue was mashed and filtered through 70 μm cell strainer with R10. For spleens, red blood cells (RBC) were lysed using RBC lysing buffer, containing ammonium chloride, potassium bicarbonate, EDTA and MiliQ water. The SI-LP cell isolation was performed as described previously(Luda et al., 2016).

### Flow cytometer

Ca/Mg-containing PBS with 2% FCS was used as buffer during the entire staining procedure. Non-specific binding was blocked with rat anti-mouse CD16/CD32 Fc block (2.4G2, BD Biosciences) for 20 minutes at 4°C. Dead cells identified as propidium iodide^+^ (Sigma Aldrich) or by Aqua LIVE/DEAD Fixable Dead Cell Staining Kit (Life Technologies) and cell aggregates (identified on FSC-A versus FSC-H scatterplots) were excluded from analyses. DCs were identified by using the following antibodies: α-CD3 (145-2C11), α-CD19 (eBio1D3), α-NK1.1 (PK136), α-B220 (RA3-6B2), α-CD64 (X54-5/7.1), α-CD103 (M290), α-CD11b (M1/70), α-CD11c (HL3), α-CD8a (53-6.7), α-CD86 (GL1), α-MHC-II I (IA/I-E) (M5/114.15.2), α-CD45.1 (A20), α-CD45.2 (104), α-IFNAR (MAR1-5A3), α-XCR1 (ZET), α-SiglecH (551), and α-TLR3 (11F8). Intracellular staining was performed using the FoxP3 Fixation/Permeabilization Kit (eBioscience) according to the manufacturer’s instructions. Data was acquired on a FACS Aria II or LSRII (BD Biosciences) and analyzed using FlowJo software (Tree Star).

### Adoptive Transfers

Bone marrow (BM) chimeras were generated by intravenous injection of BM (5 x 10^6^) cells into irradiated (9 Gy) recipients. Analysis of BM chimeras was performed 6-8 weeks after cell transfer. In all mixed BM chimeras, WT cells were identified by CD45.1 expression.

### Real-Time PCR

Total RNA was isolated from small intestine (SI) using the RNeasy kit (QIAGEN). cDNA was generated using iScript^™^ cDNA Synthesis Kit (Bio-Rad). Quantitative PCR was performed on a CFX96™ Real-Time PCR Detection System (Bio-Rad), using SsoFast^™^EvaGreen^®^ Supermix (Bio-Rad). The expression of all genes was normalized to the mean of beta-actin, Reep5 and GAPDH. Primer sequences are specified in Suppl. Table 1. Undetectable values were calculated based on the highest possible Cq +1 (=41cyles).

### Statistical Analysis

Statistics were performed using two-way ANOVA considering treatment and experiment as factors for the analysis. Wherever indicated in the figure legends, Mann-Whitney U test was applied to compare two groups (e.g.: different treatments (n=2) within the same genotype), and Kruskal-Wallis test was applied to compare more than 2 groups (e.g.: different genotypes (n=3) within the same treatment). Statistical significance was estimated by using R Studio.

R Scripts:

- Two-way ANOVA: aov(value ~ genotype + day, data = Data)

- Post-Hoc test Tukey: TukeyHSD(Data_anova2, which=“genotype”)
- Mann-Whitney U test: wilcox.test(value ~ genotype, data = Data, exact = FALSE)

Where genotype accounts for analysis of different genotypes within the same treatment. Using treatment instead of genotype allows for analysis of different treatments within a genotype. Data refers to the data to be analyzed.

## Supplementary Data

**Suppl.1:**
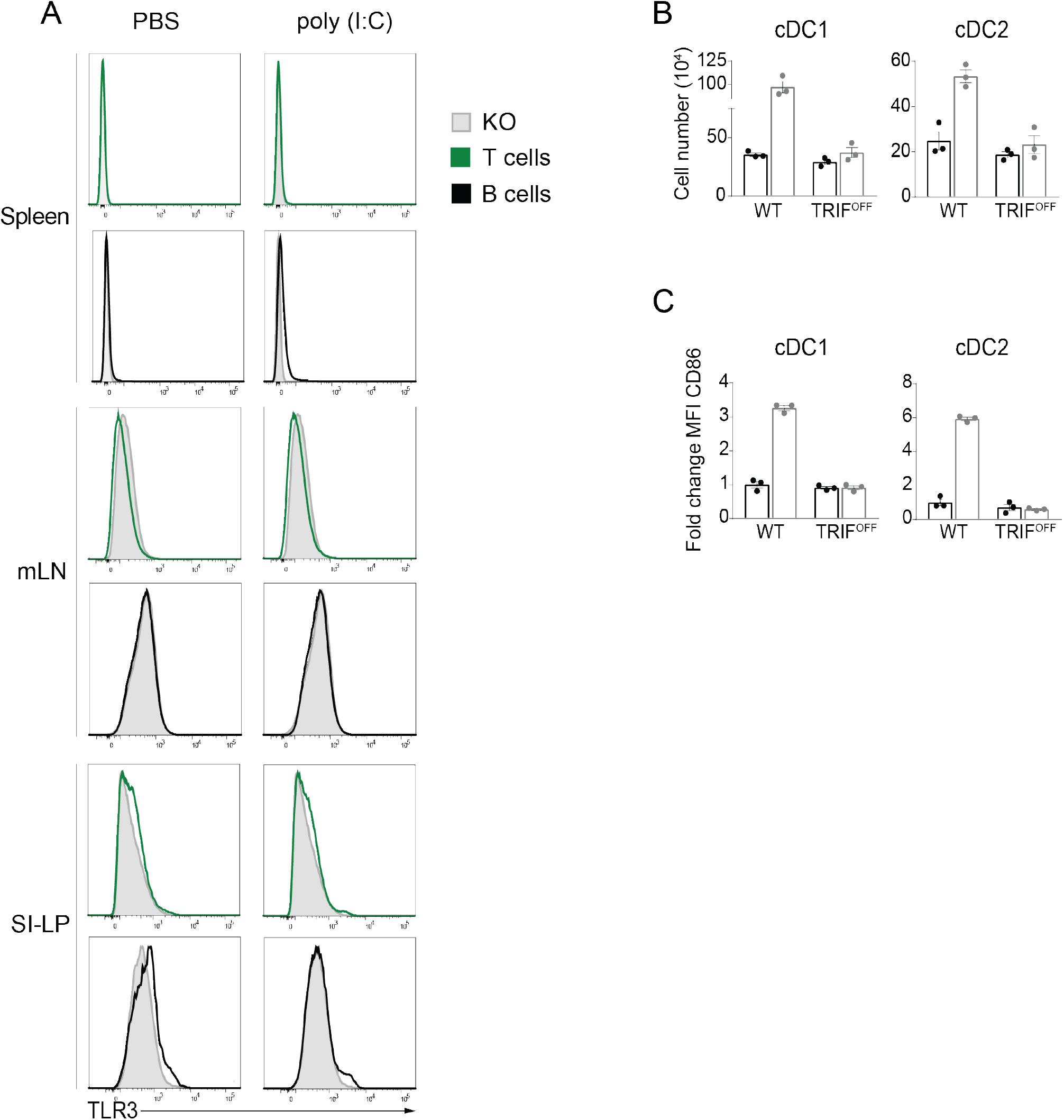
A. Intracellular TLR3 staining of B and T cells 12 hours after i.p injection of PBS or 100μg poly(I:C) into wild type mice and by B and T cells from resting TLR3^OFF^. B. Total numbers of cDC1 and cDC2 in the mLNs of WT and TRIF^OFF^ mice 12h after i.p. injection of PBS or 100μg poly(I:C). Data shown are mean numbers of cells ± 1 sem of one experiment representative of three with 3 mice per group. Mann Whitney U test, not significant. C. Activation of cDC subsets in mLNs of WT and TRIF^OFF^ mice by poly(I:C). Results shown are fold changes in CD86 expression 12h after injection of 100μg poly(I:C) as assessed by MFI normalized to FMO and relative to expression by DCs in untreated WT. Data shown are means ± 1 sem of one experiment representative of three with 3 mice per group. Mann Whitney U test, not significant.

**Suppl.2:**
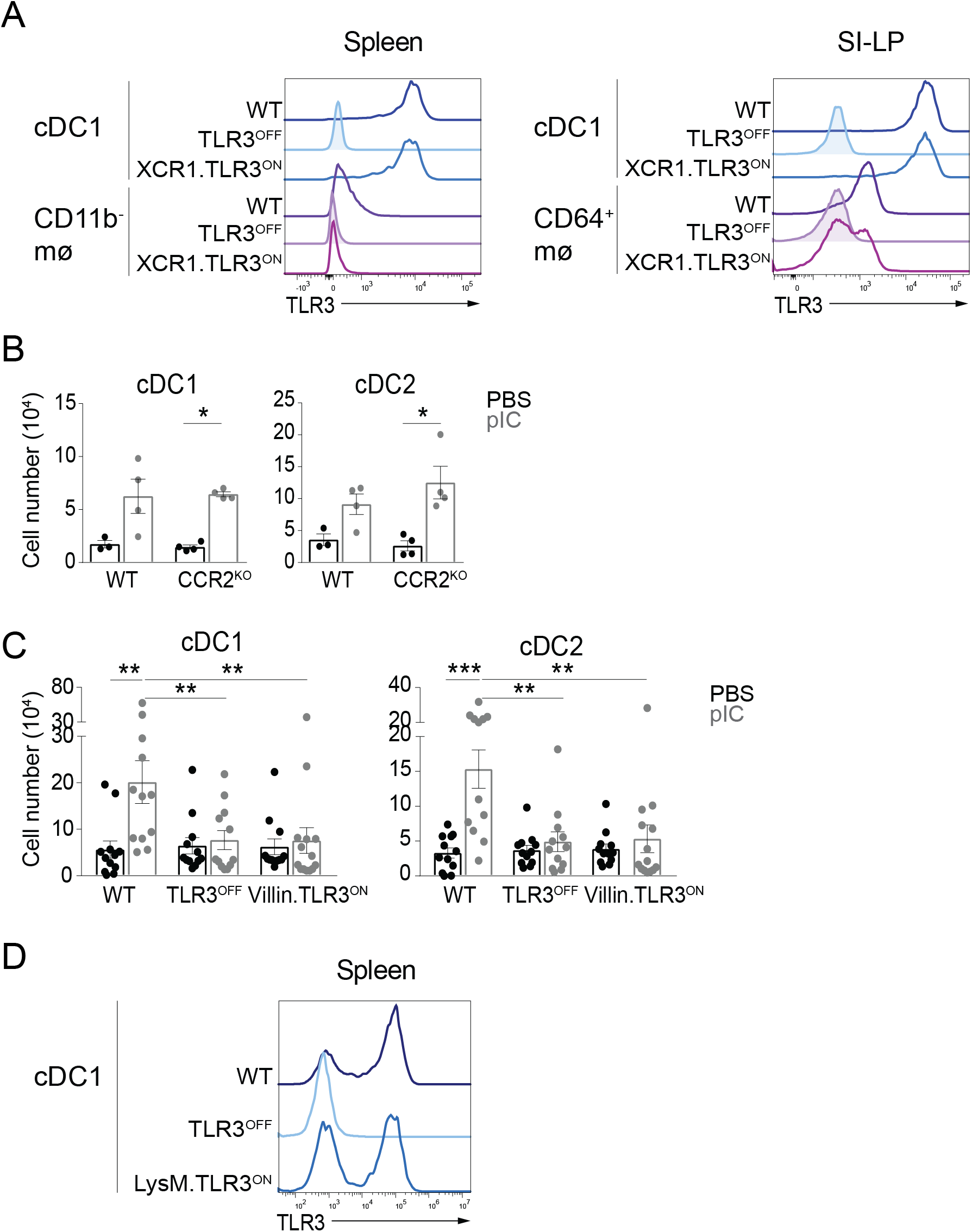
A. Intracellular TLR3 staining of spleen and SI LP in the indicated DC and macrophage populations from WT, TLR3^OFF^ and XCR1.TLR3^ON^ mice. B. Total number of cDC1 and cDC2 in the mLNs of WT and CCR2^KO^ mice 12h after i.p. injection of PBS or 100μg poly(I:C). Data shown are mean numbers of cells ± 1 sem from one experiment with 4 mice per group. Mann Whitney test, *p<0.05. C. Total number of cDC1 and cDC2 in the mLNS of WT, TLR3^OFF^ and Villin.TLR3^ON^ mice 12h after i.p. injection of PBS or 100μg poly(I:C). Data shown are mean numbers of cells ± 1 sem pooled from four independent experiments with 3 mice per group. Two-way ANOVA, **p<0.005, ***p<0.0005. D. Intracellular TLR3 staining of spleen cDC1 from WT, TLR3^OFF^ and LysM.TLR3^ON^ mice. Data shows one representative mouse per genotype from two independent experiments with 3-4 mice per group.

**Table 1:**
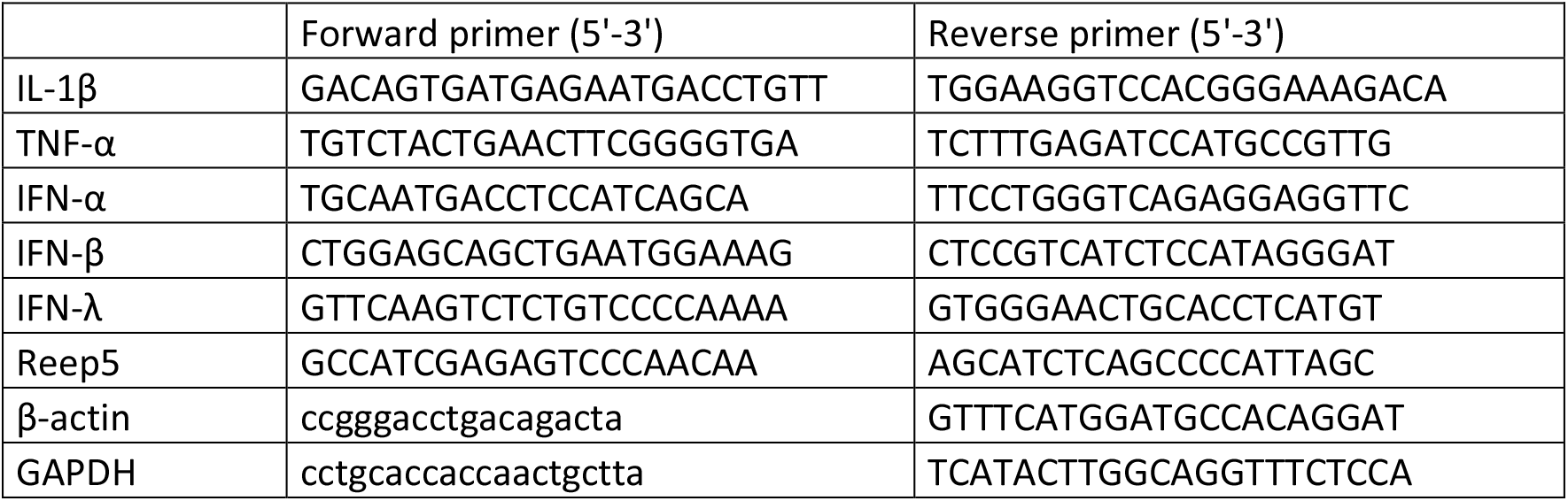
Primer Sequences for RT-PCR

